# Sexual selection and sexual size dimorphism in animals

**DOI:** 10.1101/2021.05.10.443408

**Authors:** Tim Janicke, Salomé Fromonteil

## Abstract

Sexual selection is often considered as a critical evolutionary force promoting sexual size dimorphism (SSD) in animals. However, empirical evidence for a positive relationship between sexual selection on males and male-biased SSD received mixed support depending on the studied taxonomic group and on the method used to quantify sexual selection. Here, we present a meta-analytic approach accounting for phylogenetic non-independence to test how standardized metrics of the opportunity and strength of pre-copulatory sexual selection relate to SSD across a broad range of animal taxa comprising up to 102 effect sizes from 64 species. We found that SSD was correlated with the sex difference in the opportunity for sexual selection but not with the sex difference in the Bateman gradient. These findings suggest that pre-copulatory sexual selection plays a limited role for the evolution of sexual size dimorphism in a broad phylogenetic context.

## Introduction

Females and males often show striking differences in morphological, behavioural, and physiological traits. Understanding the evolutionary drivers for this diversity is at the heart of sexual selection research (Andersson 1994). One of the best studied sex differences is dimorphism in body size, which can be found in animals along a broad continuous spectrum ranging from extremely females-biased (e.g. spiders; (Foellmer & Moya-Larano 2007)) towards extremely male-biased dimorphism (e.g. sea elephants; (Lindenfors *et al*. 2002)). Theory predicts that any sex difference such as sexual size dimorphism (hereafter SSD) is ultimately rooted in anisogamy and the resulting sexual selection (Schärer *et al*. 2012), defined as selection arising from competition for mating partners and/or their gametes (Shuker 2010). Specifically, body size is often assumed to be sexually selected because it may confer an advantage during pre- and/or post-copulatory mate competition and mate choice. Such sexual selection on body size may shift the evolutionary optimum of the sex under stronger competition away from the optimum from the other sex, which manifests in intra-locus sexual conflict and can eventually be resolved in terms of sexual dimorphism (Lande 1980; Reeve & Fairbairn 2001). However, the primordial effects of anisogamy and sexual selection on the evolution of sexual dimorphism can be altered by sex-specific selection on traits that affect the competition for resources other than mates. This so-called ecological character displacement has recently been argued to be key for complete understanding of sexual dimorphism in a variety of traits, including feeding morphology, habitat use and especially body size (De Lisle 2019). Therefore, both sexual and natural selection may contribute to the evolution of SSD and potentially in opposing ways (Fairbairn *et al*. 2007).

There is compelling evidence from numerous comparative studies suggesting that sexual selection plays indeed an important role for the evolution of SSD in many animal taxa including insects (Serrano-Meneses *et al*. 2008), fishes (Pyron *et al*. 2013), birds (Szekely *et al*. 2007), reptiles (Shine 1994), and mammals (Cassini 2020). However, for many of those taxa there are also comparative studies that failed to demonstrate a relationship between sexual selection and SSD (Figuerola & Green 2000; Kratochvil & Frynta 2002; Pincheira-Donoso *et al*. 2021) or conclude that sexual selection plays only a minor role in shaping size difference between males and females (Cox *et al*. 2003). These inconsistencies may arise from differences within tested clades but may also stem from difficulties in quantifying the strength and direction of sexual selection in a meaningful way allowing inter-specific comparisons. In fact, most studies use categorical variables to describe inter-specific variation in sexual selection such as mating system or male combat, which often do not capture the entire variation in the strength of sexual selection. Moreover, many studies use proxies that can only measure sexual selection in males (e.g. of harem size, testis size, male territoriality) assuming that competition for mates is absent or rare in females, which might often be an oversimplification (Hare & Simmons 2019).

Here, we aim to expand our knowledge on the evolution of sexual size dimorphism by testing its relationship with standardised metrics of sex-specific sexual selection derived from Bateman’s principles (Bateman 1948). Specifically, we use a phylogenetic meta-analysis to explore how SSD relates to sex differences in both the opportunity for sexual selection and the Bateman gradient across a broad range of animal taxa. The opportunity for sexual selection (*I*_s_) is defined as the variance in relativized mating success, which provides a proxy of the intensity of competition for mating partners in a population (Jones 2009). The Bateman gradient (*β*_ss_) represents the slope of a linear regression of reproductive success on mating success and provides an upper limit of pre-copulatory sexual selection on a phenotypic trait (Arnold 1994). Both metrics have been argued to provide reliable and complementary measures for the strength of sexual selection (Mobley 2014; Anthes *et al*. 2017) of which *β*_ss_ quantifies the benefit of having an additional mating partner and *I*_s_ the difficulty in obtaining such an additional mate (Kokko *et al*. 2006). If sexual selection drives sexual size dimorphism, we expect the sex with the steeper *β*_ss_ and the higher *I*_s_ to be the bigger relative to the other.

## Methods

### Data acquisition

We extracted published estimates of the strength of sexual selection following a search protocol described in more detail elsewhere (Janicke *et al*. 2016). In brief, we conducted a systematic literature search using the ISI Web of Knowledge (Web of Science Core Collection, from 1900 to 2020) with the ‘topic’ search terms defined as (‘Bateman*’ OR ‘opportunit* for selection’ OR ‘opportunit* for sexual selection’ OR ‘selection gradient*’). All primary articles were screened for studies reporting both male and female estimates for *I*_s_ or *β*_ss_. We used a database updated on the 30^th^of November 2020 (for PRISMA diagram see Figure S1) encompassing 93 estimates of *I*_s_ and 102 estimates of *β*_ss_ from 71 studies.

In order to estimate SSD for all sampled species, we extracted estimates of male and female size from the literature (Table S1). Whenever possible we used linear length measures such as total body length (*N* = 25), wing length (*N* = 9), snout-to-vent length (*N* = 4), elytra length (*N* = 4) and thorax length (*N* = 3). For species for which we could not find length measures we used body weight (*N* = 14) as a proxy for size. We then computed sexual size dimorphism as

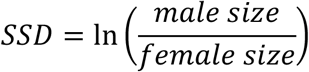

with positive or negative values indicating species with males or females being the larger sex, respectively.

### Phylogenetic affinities

To account for phylogenetic nonindependence, we reconstructed the phylogeny of all sampled species using divergence times from the TimeTree database (http://www.timetree.org/; (Kumar *et al*. 2017)). The obtained distance matrix was then transformed into the NEWICK format using the unweighted pair group method with arithmetic mean (UPGMA) algorithm implemented in MEGA (https://www.megasoftware.net/; (Kumar *et al*. 2018)). In total, our dataset included 64 species from a broad range of animal taxa with a majority of arthropods (*N*_Species_ = 16), birds (*N*_Species_ = 13), fishes (*N*_Species_ = 8), and mammals (*N*_Species_ = 8) (Figure S2).

### Statistical analysis

We computed effect sizes of the sex difference in *I*_s_ or *β*_ss_ as described in Janicke *et al*. (2016). In particular, we used the natural logarithm of the ratio between male and female coefficients of variation in mating success (ln*CVR*; (Nakagawa et al. 2015)) as an effect size for the sex difference in the opportunity for sexual selection (Δ*I*_s_). Moreover, we computed Hedges’ *g* (Borenstein et al. 2009) by subtracting the female from the male Bateman gradient, in order to obtain an effect size for the sex difference in in the Bateman gradient (Δ*β*_ss_). Hence, positive values of Δ*I*_s_ and Δ*β*_ss_ indicate a male bias in the opportunity and the strength of sexual selection, respectively.

We tested for a relationship between SSD and sex-specific sexual selection by running Generalized Linear Mixed Effects Models (GLMMs) using the rma.mv function implemented in the metafor R package version 2.4-0 (Viechtbauer 2010). In particular, we defined the effect size (Δ*I*_s_ or Δ*β*_ss_) as response variable weighted by the inverse of its sampling variance and included SSD as the only fixed predictor variable. In all models we included study identifier, observation identifier and species identifier (phylogenetic correlation matrix) as random terms. Goodness of fit was quantified using McFadden’s *R*^2^ inferred from the ratio of the log-likelihoods of the full model (with SSD as predictor variable) and the null model (no predictor variable included) (Sokal & Rohlf 2012).

Outlier analysis of estimates of SSD revealed that one of the sampled species (*Latrodectus hasseltii*) has an extremely female biased SSD (Grubb’s test: *G* = 5.760, *P* < 0.001) and was therefore excluded from the analysis. Moreover, five species are simultaneous hermaphrodites, which by definition cannot exhibit sexual dimorphism in size. For completeness, we report alternative analyses including the outlier species or excluding simultaneous hermaphrodites in the Supplementary Material (Table S2) indicating that results are robust.

We used Kendell’s rank correlation test to quantify funnel plot asymmetry, which can be indicative of publication bias. Residuals of all models were checked for normality by visual inspection of quantile-quantile plots. All statistical analyses were carried out in R version 4.0.3 (R Core Team 2020).

## Results

We found evidence for a positive relationship between SSD and Δ*I*_s_ (phylogenetic GLMM: *N* = 93, Estimate ± SE = 1.014 ± 0.324, *Q*_M_ = 9.792, *P* = 0.002, McFadden’s *R*^2^ = 0.09) with an intercept not differing significantly from zero (intercept ± SE = 0.170 ± 0.152, *z* = 1.115, *P* = 0.265). By contrast, SSD did not predict variation in Δ*β*_ss_ (phylogenetic GLMM: *N* = 101, Estimate ± SE = 0.3442 ± 0.0.341, *Q*_M_ = 1.019, *P* = 0.313, McFadden’s *R*^2^ = 0.01) with a significantly positive intercept (intercept ± SE = 0.397 ± 0.058, *z* = 6.801, *P* < 0.001).

We detected signs for funnel plot asymmetry (Figure S3) for Δ*I*_s_ (Kendall’s tau = 0.274, *P* < 0.001) and Δ*β*_ss_ (Kendall’s tau = 0.222, *P* < 0.001) with an underrepresentation of studies of low precision and a negative effect-size (i.e., stronger sexual selection in females).

## Discussion

This study provides, to our knowledge the first test for an evolutionary link between standardized metrics of the strength of sexual selection and SSD across a broad phylogenetic range covering vertebrate and non-vertebrate taxa. We found mixed support for the hypothesis that sexual selection is a main driver for the evolution of sexual size dimorphism. Specifically, SSD was correlated with the Δ*I*_s_ but not Δ*β*_ss_. As predicted, species with male-biased SSD showed a higher *I*_s_ in males relative to females. This suggests that a highly skewed mating success towards bigger males may indeed promote the evolution of bigger males. Interestingly, our statistical model predicted species in which sexes do not differ in size to show no sex-bias in *I*_s_, as indicated by a statistically non-significant intercept. This bolsters the idea that a sex difference in *I*_s_ often translates into a sex difference in size. However, the observed effect was weak with SSD explaining only 9% of the variance in Δ*I*_s_ (equivalent to an effect size of *r* = 0.3). In a previous study, Soulsbury *et al*. (2014) tested for a relationship between the *I*_s_ and SSD reporting evidence for a relationship in mammals but not in birds. However, their analysis was loaded primarily by estimates of the opportunity for selection (i.e., variance in reproductive success; *I*) rather than *I*_s_, which makes their findings difficult to interpret especially because *I* does not only capture the opportunity for sexual but also for natural selection (Mobley 2014). Moreover, their study included only estimates of *I* and *I*_s_ of males assuming that sexual selection on body size can be neglected in females, which might often be an incorrect assumption (Hare & Simmons 2019).

Contrary to our findings on Δ*I*_s_, we did not detect a relationship between SSD and Δ*β*_ss_ suggesting that sex-specific selection on body size is not mediated by sex-differences in pre-copulatory sexual selection. These contrasting results may reflect differences in the properties of *I*_s_ and *β*_ss_, with the former providing a measure of the potential for intra-sexual competition for mates whereas the latter quantifies the benefit of succeeding in mate acquisition (Kokko *et al*. 2006). Nevertheless, given that *β*_ss_ constitutes a proxy for the upper potential of pre-copulatory sexual selection on body size (Jones 2009), we would have predicted an effect on Δ*β*_ss_ if sexual selection was a strong determinant of SSD in animals.

Both proxies of sexual selection used in this study have been argued to be particularly powerful in comparative studies (Mobley 2014; Anthes *et al*. 2017) but like other proxies they also have their limitations. Notably, *I*_s_ and *β*_ss_ measure primarily the potential and strength of pre-copulatory sexual selection. This might be especially problematic for the male sex, in which sexual selection on body size can operate also along post-copulatory episodes, because bigger individuals can be better sperm competitors (Pélissié *et al*. 2014). Presumably, the most powerful approach to test for an evolutionary link between sexual selection and SSD across species involves mating- and post-mating differentials of body size for both sexes (Jones 2009). We presume that this kind of data is currently available for only a few species but might become available in the future to be subjected to a broad scale meta-analysis.

Collectively, our study provides limited evidence for the hypothesis that sexual selection promotes SSD when tested across a broad phylogenetic context. Despite our finding of a positive relationship between Δ*I*_s_ and SSD, its explanatory power was low suggesting that sexual selection plays only a minor role for the evolution of SSD. Moreover, our results echo the body of previous comparative studies indicating that evidence for an effect of sexual selection and SSD depends on the proxy used to quantify sexual selection. For these reasons, our study challenges the common practice of using SSD as a proxy for sexual selection (Chen *et al*. 2012; Kahrl *et al*. 2016; Mikula *et al*. 2021). We conclude that despite compelling evidence for SSD to correlate with sexual selection within some animal clades, it appears to be a poor predictor for pre-copulatory sexual selection at a broader taxonomic scale. Alternative mechanisms such as ecological character displacement may be crucial to understand the full diversity of SSD in animals.

## Acknowledgements

We are grateful to the authors of all primary studies for making their research accessible. This study was funded by the German Research Foundation (DFG grant number: JA 2653/2-1) and the CNRS.

## Figure Legends

**Figure 1.**
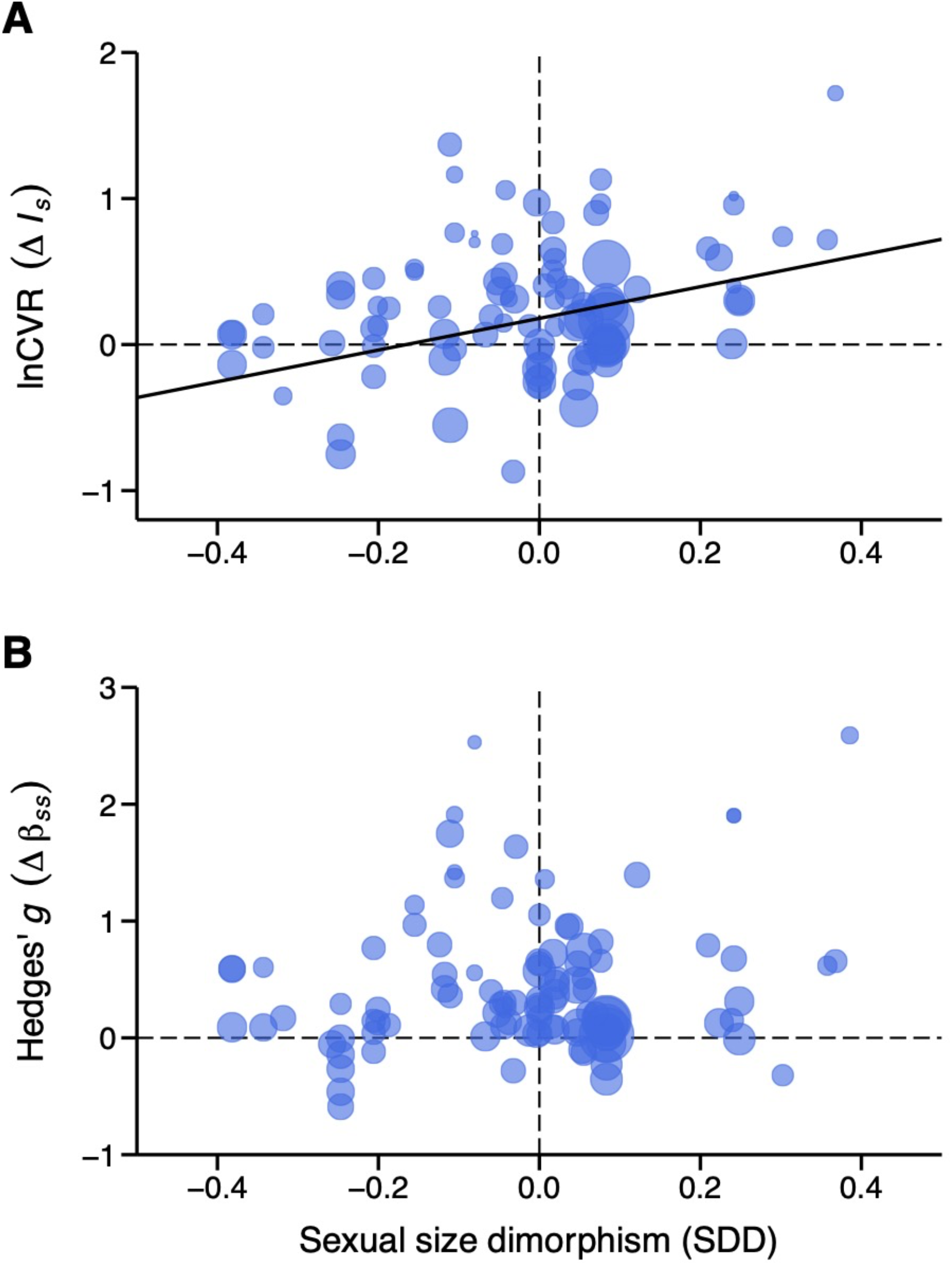
Relationship between sexual size dimorphism (SDD) and sexual selection quantified in terms of (A) the sex difference in the opportunity for sexual selection (Δ*I*_s_) and (B) the sex difference in the Bateman gradient (Δ*β*_ss_). Positive values in SDD, Δ*I*_s_ and Δ*β*_ss_ indicate a male bias.

