## Supplementary Figures and Tables for "Sexual selection and sexual size dimorphism in animals"

**Contents**

Figure S1 – S3. (pp. 2 – 4)

Table S1 – S2. (pp. 5 – 10)

Supplementary Data: List of primary studies (pp. 11 – 15)

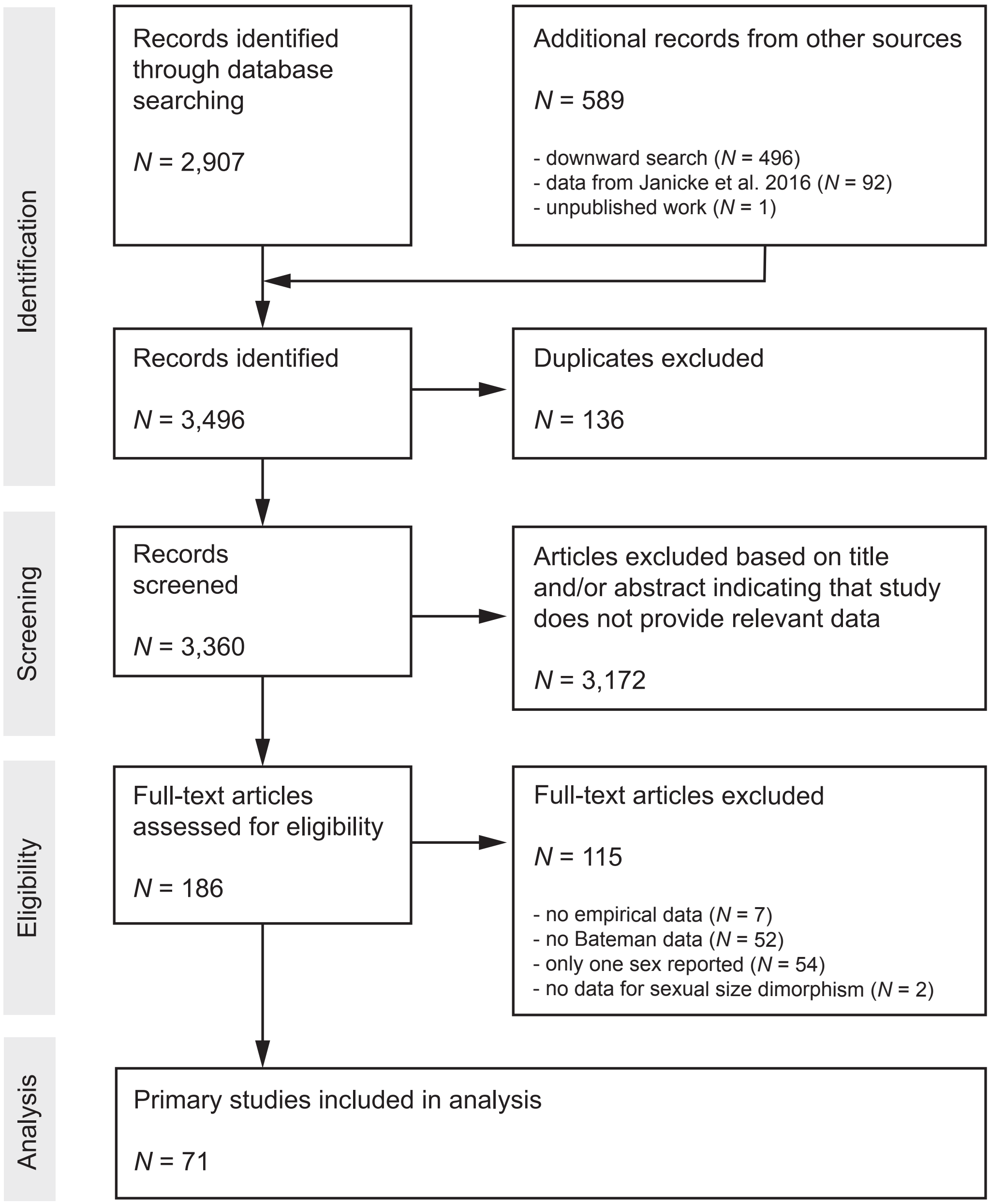


**Figure S1. Preferred Reporting Items for Systematic Reviews and Meta-Analyses (PRISMA) Diagram.** Flow chart shows the number of records identified during the different phases of the systematic literature search.

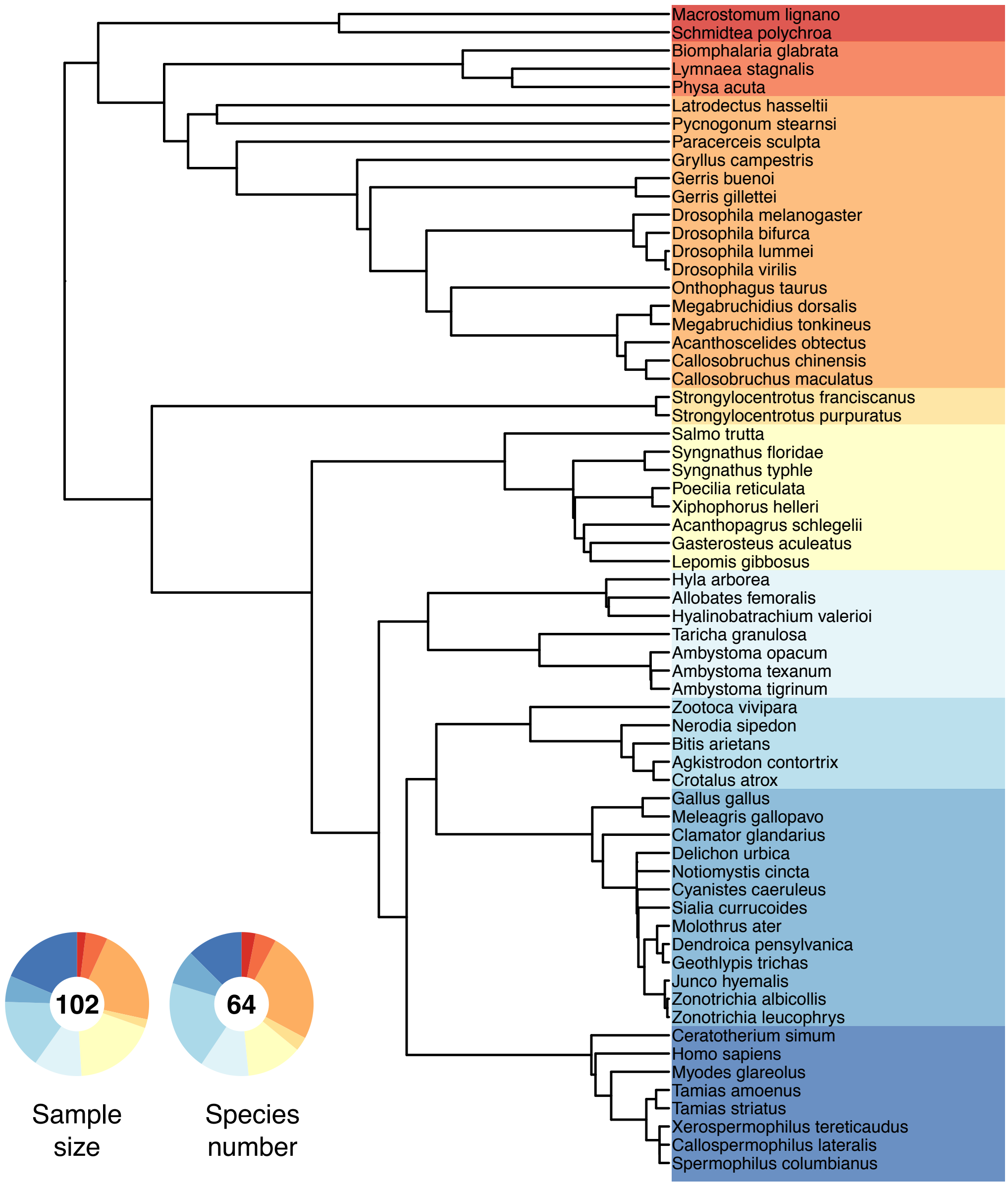


**Figure S2. Phylogenetic tree of all sampled species.** Doughnut charts show the relative fraction of the sampled effect sizes (i.e., Hedges's *g* of $\Delta\beta_{ss}$) and the number of species.

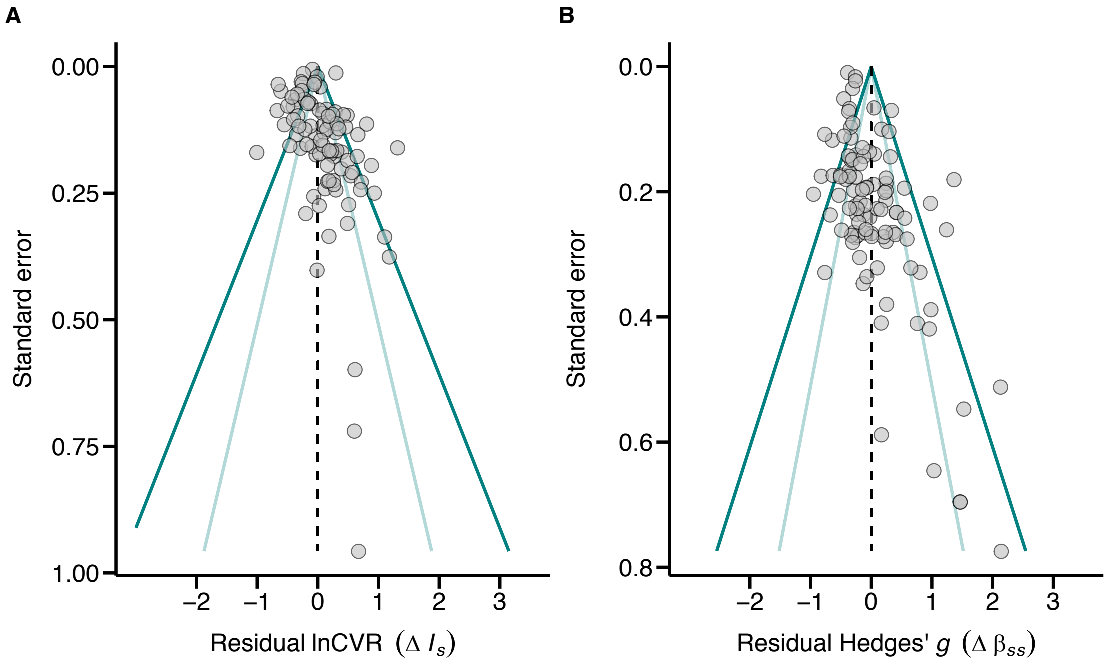


**Figure S3. Funnel plots for lnCVR ($\Delta l_s$) and Hedges'g ($\Delta \beta_{ss}$).** Data are shown for meta-analytic residuals obtained from multivariate linear mixed-effects models accounting for random terms (i.e., phylogeny, species, study identifier) and sexual size dimorphism. Dashed lines indicate the estimated global effect size. Dark and light blue solid lines denote the expected 95% and 99% confidence limits purely due to sampling heterogeneity. Asymmetries along the global effect size may reflect publication biases.

**Table S1.** Estimates of sexual size dimorphism (SSD) with references for all sampled species. SSD was estimated as ln(male size/female size) so that positive or negative values indicate species with males or females being the larger sex, respectively.

| Class | Species | SSD | Reference |
|---|---|---|---|
| Mammalia | *Callospermophilus lateralis* | -0.029 | (Iwaniuk 2001) |
| | *Ceratotherium simum* | 0.302 | (MacDonald 2010) |
| | *Homo sapiens* | 0.083 | (McDowell *et al.* 2008) |
| | *Myodes glareolus* | -0.105 | (Bondrup-Nielsen & Ims 1990) |
| | *Spermophilus columbianus* | 0.121 | (Dobson 1992) |
| | *Tamias amoenus* | -0.124 | (Schulte-Hostedde & Millar 2000) |
| | *Tamias striatus* | -0.012 | (Forbes 1966) |
| | *Xerospermophilus tereticaudus* | 0.022 | (Munroe & Koprowski 2011) |
| Reptilia | *Agkistrodon contortrix* | 0.210 | (Smith *et al.* 2008) |
| | *Bitis arietans* | 0.071 | (Shine 1994) |
| | *Crotalus atrox* | 0.060 | (Beaupre *et al.* 1998) |
| | *Nerodia sipedon* | -0.111 | (Weatherhead *et al.* 1995) |
| | *Zootoca vivipara* | -0.081 | (Massot *et al.* 1992) |
| Aves | *Clamator glandarius* | 0.053 | (Macias-Sanchez *et al.* 2013) |
| | *Cyanistes caeruleus* | 0.047 | (Blondel *et al.* 2002) |
| | *Delichon urbica* | 0.007 | (de Lope 1986) |
| | *Dendroica pensylvanica* | 0.035 | (Kissner *et al.* 2003) |
| | *Gallus gallus* | 0.358 | (Parker & Garant 2005) |
| | *Geothlypis trichas* | 0.054 | (Morris *et al.* 2003) |
| | *Junco hyemalis* | 0.055 | (Nolan & Ketterson 1983) |
| | *Meleagris gallopavo* | 0.368 | (Ajayi *et al.* 2012) |
| | *Molothrus ater* | 0.248 | (Johnson *et al.* 1980) |
| | *Notiomystis cincta* | 0.223 | (Walker *et al.* 2014) |
| | *Sialia currucoides* | 0.038 | (Billerman *et al.* 2020) |
| | *Zonotrichia albicollis* | 0.049 | (Schlinger 1990) |
| | *Zonotrichia leucophrys* | 0.048 | (Mewaldt & King 1986) |
| Amphibia | *Allobates femoralis* | -0.038 | (Ursprung *et al.* 2011) |
| | *Ambystoma opacum* | -0.155 | (Pokhrel 2009) |
| | *Ambystoma texanum* | -0.318 | (Williams *et al.* 2009) |
| | *Ambystoma tigrinum* | -0.343 | (Williams *et al.* 2009) |
| | *Hyalinobatrachium valerioi* | -0.053 | (Mangold *et al.* 2015) |
| | *Hyla arborea* | -0.042 | (Ozdemir *et al.* 2012) |
| | *Taricha granulosa* | 0.242 | (Janzen & Brodie 1989) |
| Actinopterygii | *Acanthopagrus schlegelii* | -0.187 | (Horne *et al.* 2020) |
| | *Gasterosteus aculeatus* | -0.201 | (Horne *et al.* 2020) |
| | *Lepomis gibbosus* | -0.003 | (Jastrebski 2001) |
| | *Poecilia reticulata* | -0.382 | (Bisazza & Pilastro 1997) |

| Class | Species | SSD | Reference |
|---|---|---|---|
| | *Salmo trutta* | 0.019 | (Petersson & Jarvi 1997) |
| | *Syngnathus floridae* | -0.032 | (Mobley *et al.* 2014) |
| | *Syngnathus typhle* | -0.247 | (Rispoli & Wilson 2008) |
| | *Xiphophorus helleri* | 0.076 | (Basolo & Wagner 2004) |
| Echinoidea | *Strongylocentrotus franciscanus* | 0.000 | (Conor 1972) |
| | *Strongylocentrotus purpuratus* | 0.000 | (Conor 1972) |
| Arthropoda | *Acanthoscelides obtectus* | -0.206 | (Seslija *et al.* 2009) |
| | *Callosobruchus chinensis* | -0.060 | (Colgoni & Vamosi 2006) |
| | *Callosobruchus maculatus* | -0.048 | (Colgoni & Vamosi 2006) |
| | *Drosophila bifurca* | -0.031 | (Pitnick *et al.* 1995) |
| | *Drosophila lummei* | -0.044 | (Pitnick *et al.* 1995) |
| | *Drosophila melanogaster* | -0.118 | (Pitnick *et al.* 1995) |
| | *Drosophila virilis* | -0.046 | (Pitnick *et al.* 1995) |
| | *Gerris buenoi* | -0.111 | (Andersen 1996) |
| | *Gerris gillettei* | -0.067 | (Arnqvist 1997) |
| | *Gryllus campestris* | -0.044 | (Harz 1969) |
| | *Latrodectus hasseltii* | -1.189 | (Andrade 1996) |
| | *Megabruchidius dorsalis* | 0.239 | (Takakura 1999) |
| | *Megabruchidius tonkineus* | 0.056 | (Salehialavi *et al.* 2011) |
| | *Onthophagus taurus* | 0.017 | (Palestrini *et al.* 2000) |
| | *Paracerceis sculpta* | 0.386 | (Shuster & Guthrie 1999) |
| | *Pycnogonum stearnsi* | -0.257 | (Barreto & Avise 2010) |
| Gastropoda | *Biomphalaria glabrata* | 0.000 | - [1] |
| | *Lymnaea stagnalis* | 0.000 | - [1] |
| | *Physa acuta* | 0.000 | - [1] |
| Turbellaria | *Macrostomum lignano* | 0.000 | - [1] |
| | *Schmidtea polychroa* | 0.000 | - [1] |

[1] Simultaneous hermaphrodite in which both sexes expressed in the same individual.

#### References – Estimation of sexual size dimorphism


Ajayi, O.O., Yakubu, A., Jayeola, O.O., Imumorin, I.G., Takeet, M.I., Ozoje, M.O. *et al.* (2012). Multivariate analysis of sexual size dimorphism in local turkeys (*Meleagris gallopavo*) in Nigeria. *Trop. Anim. Health Prod.*, 44, 1089-1095.

Andersen, N.M. (1996). Ecological phylogenetics of mating systems and sexual dimorphism in water striders (Heteroptera: Gerridae). *Vie Milieu*, 46, 103-114.

Andrade, M.C.B. (1996). Sexual selection for male sacrifice in the Australian redback spider. *Science*, 271, 70-72.

Arnqvist, G. (1997). The evolution of water strider mating systems: causes and consequences of sexual conflicts. In: *The evolution of mating systems in insects and arachnids* (eds. Choe, JC & Crespi, BJ). Cambridge University Press Cambridge, UK, pp. 146-163.

Barreto, F.S. & Avise, J.C. (2010). Quantitative measures of sexual selection reveal no evidence for sex-role reversal in a sea spider with prolonged paternal care. *Proc. R. Soc. B-Biol. Sci.*, 277, 2951-2956.

Basolo, A.L. & Wagner, W.E. (2004). Covariation between predation risk, body size and fin elaboration in the green swordtail, *Xiphophorus helleri*. *Biol. J. Linnean Soc.*, 83, 87-100.

Beaupre, S.J., Duvall, D. & O'Leile, J. (1998). Ontogenetic variation in growth and sexual size dimorphism in a central Arizona population of the western diamondback rattlesnake (*Crotalus atrox*). *Copeia*, 40-47.

Billerman, S.M., Keeney, B.K., Rodewald, P.G. & Schulenberg, T.S. (2020). Birds of the World. Available at: https://birdsoftheworld.org/bow/.

Bisazza, A. & Pilastro, A. (1997). Small male mating advantage and reversed size dimorphism in poeciliid fishes. *J. Fish Biol.*, 50, 397-406.

Blondel, J., Perret, P., Anstett, M.C. & Thebaud, C. (2002). Evolution of sexual size dimorphism in birds: test of hypotheses using blue tits in contrasted Mediterranean habitats. *J. Evol. Biol.*, 15, 440-450.

Bondrup-Nielsen, S. & Ims, R.A. (1990). Reversed sexual size dimorphism in microtines: Are females larger than males or are males smaller than females? *Evol. Ecol.*, 4, 261-272.

Colgoni, A. & Vamosi, S.M. (2006). Sexual dimorphism and allometry in two seed beetles (Coleoptera : Bruchidae). *Entomol. Sci.*, 9, 171-179.

Conor, J. (1972). Gonad growth in the sea urchin, *Strongylocentrotus purpuratus* (Stimpson) (Echinodermata: Echinoidea) and the assumptions of gonad index methods. *J. Exp. Mar. Biol. Ecol.*, 10, 89-103.

de Lope, F. (1986). The biometry of House Martin (*Delichon urbica* L.). *Ardeola*, 33, 171-201.

Dobson, F.S. (1992). Body mass, structural size, and life-history patterns of the columbian ground squirrel. *Am. Nat.*, 140, 109-125.

Forbes, R.B. (1966). Studies of the biology of Minnesotan chipmunks. *American Midland Naturalist*, 76, 290-308.

Harz, K. (1969). *The Orthoptera of Europe*. Springer, The Hague.

Horne, C.R., Hirst, A.G. & Atkinson, D. (2020). Selection for increased male size predicts variation in sexual size dimorphism among fish species. *Proc. R. Soc. B-Biol. Sci.*, 287, 9.

Iwaniuk, A.N. (2001). Interspecific variation in sexual dimorphism in brain size in Nearctic ground squirrels (*Spermophilus* spp.). *Can. J. Zool.*, 79, 759-765.

Janzen, F.J. & Brodie, E.D. (1989). Tall tails and sexy males: sexual behavior of rough-skinned newts (*Taricha granulosa*) in a natural breeding pond. *Copeia*, 1989, 1068-1071.

Jastrebski, C.J. (2001). Divergence and selection in trophically polymorphic pumpkinseed sunfish *Lepomis gibbosus*. University of Guelph, p. 121.

Johnson, D.M., Stewart, G.L., Corley, M., Ghrist, R., Hagner, J., Ketterer, A. *et al.* (1980). Brown-Headed Cowbird (*Molothrus ater*) Mortality in an Urban Winter Roost *Auk*, 97, 299-320.

Kissner, K.J., Weatherhead, P.J. & Francis, C.M. (2003). Sexual size dimorphism and timing of spring migration in birds. *J. Evol. Biol.*, 16, 154-162.

MacDonald, D.W. (2010). *The encyclopedia of mammals*. Oxford University Press.

Macias-Sanchez, E., Martinez, J.G., Aviles, J.M. & Soler, M. (2013). Sexual differences in colour and size in the Great Spotted Cuckoo *Clamator glandarius*. *Ibis*, 155, 605-610.

Mangold, A., Trenkwalder, K., Ringler, M., Hodl, W. & Ringler, E. (2015). Low reproductive skew despite high male-biased operational sex ratio in a glass frog with paternal care. *BMC Evol. Biol.*, 15, 13.

Massot, M., Clobert, J., Pilorge, T., Lecomte, J. & Barbault, R. (1992). Density dependence in the common lizard: Demographic consequences of a density manipulation. *Ecology*, 73, 1742-1756.

McDowell, M.A., Fryar, C.D., Ogden, C.L. & Flegal, K.M. (2008). Anthropometric reference data for children and adults: United States, 2003–2006. *National health statistics reports*, 10.

Mewaldt, L.R. & King, J.R. (1986). Estimation of sex ratio from wing-length in birds when sexes differ in size but not coloration. *J. Field Ornithol.*, 57, 155-167.

Mobley, K.B., Abou Chakra, M. & Jones, A.G. (2014). No evidence for size-assortative mating in the wild despite mutual mate choice in sex-role-reversed pipefishes. *Ecol. Evol.*, 4, 67-78.

Morris, S.R., Donovan, A.J., Agugliaro, S.M. & Holmes, D.W. (2003). Accuracy of sex determination of hatch-year Common Yellowthroats (*Geothlypis trichas*) during the fall. *North American Bird Bander*, 28, 105-110.

Munroe, K.E. & Koprowski, J.L. (2011). Sociality, Bateman's gradients, and the polygynandrous genetic mating system of round-tailed ground squirrels (*Xerospermophilus tereticaudus*). *Behav. Ecol. Sociobiol.*, 65, 1811-1824.

Nolan, V. & Ketterson, E.D. (1983). An analysis of body mass, wing length, and visible fat deposits of Dark-Eyed Juncos wintering at different latitudes. *Wilson Bull.*, 95, 603-620.

Ozdemir, N., Altunisik, A., Ergul, T., Gul, S., Tosunoglu, M., Cadeddu, G. *et al.* (2012). Variation in body size and age structure among three Turkish populations of the treefrog *Hyla arborea*. *Amphib. Reptil.*, 33, 25-35.

Palestrini, C., Rolando, A. & Laiolo, P. (2000). Allometric relationships and character evolution in *Onthophagus taurus* (Coleoptera : Scarabaeidae). *Can. J. Zool.*, 78, 1199-1206.

Parker, T.H. & Garant, D. (2005). Quantitative genetics of ontogeny of sexual dimorphism in red junglefowl (*Gallus gallus*). *Heredity*, 95, 401-407.

Petersson, E. & Jarvi, T. (1997). Reproductive behaviour of sea trout (*Salmo trutta*) - The consequences of sea-ranching. *Behaviour*, 134, 1-22.

Pitnick, S., Markow, T.A. & Spicer, G.S. (1995). Delayed male maturity is a cost of producing large sperm in *Drosophila*. *Proc. Natl. Acad. Sci. U. S. A.*, 92, 10614-10618.

Pokhrel, L.R. (2009). Mapping the Dorsal Skin Pigmentation Patterns of Two Sympatric Populations of Ambystomatid Salamanders, *Ambystoma opacum* and *A. maculatum* from Northeast Tennessee. In: *Department of Biological Sciences*. East Tennessee State University, p. 64.

Rispoli, V.F. & Wilson, A.B. (2008). Sexual size dimorphism predicts the frequency of multiple mating in the sex-role reversed pipefish *Syngnathus typhle*. *J. Evol. Biol.*, 21, 30-38.

Salehialavi, Y., Fritzsche, K. & Arnqvist, G. (2011). The cost of mating and mutual mate choice in 2 role-reversed honey locust beetles. *Behav. Ecol.*, 22, 1104-1113.

Schlinger, B.A. (1990). A nonparametric aid in identifying sex of cryptically dimorphic birds. *Wilson Bull.*, 102, 454-550.

Schulte-Hostedde, A.I. & Millar, J.S. (2000). Measuring sexual size dimorphism in the yellow-pine chipmunk (*Tamias amoenus*). *Can. J. Zool.*, 78, 728-733.

Seslija, D., Stojkovic, B., Tucic, B. & Tucic, N. (2009). Egg-dumping behaviour in the seed beetle *Acanthoscelides obtectus* (Coleoptera: Chrysomelidae: Bruchinae) selected for early and late reproduction. *Eur. J. Entomol.*, 106, 557-563.

Shine, R. (1994). Size dimorphism in snakes revisited. *Copeia*, 326-346.

Shuster, S.M. & Guthrie, E.E. (1999). Effects of temperature and food availability on adult body length in natural and laboratory populations of *Paracerceis sculpta* (Holmes), a Gulf of California isopod. *J. Exp. Mar. Biol. Ecol.*, 233, 269-284.

Smith, C.F., Schwenk, K., Earley, R.L. & Schuett, G.W. (2008). Sexual size dimorphism of the tongue in a North American pitviper. *J. Zool.*, 274, 367-374.

Takakura, K. (1999). Active female courtship behavior and male nutritional contribution to female fecundity in *Bruchidius dorsalis* (Fahraeus) (Coleoptera : Bruchidae). *Res. Popul. Ecol.*, 41, 269-273.

Ursprung, E., Ringler, M., Jehle, R. & Hodl, W. (2011). Strong male/male competition allows for nonchoosy females: high levels of polygynandry in a territorial frog with paternal care. *Mol. Ecol.*, 20, 1759-1771.

Walker, L.K., Ewen, J.G., Brekke, P. & Kilner, R.M. (2014). Sexually selected dichromatism in the hihi *Notiomystis cincta*: multiple colours for multiple receivers. *J. Evol. Biol.*, 27, 1522-1535.

Weatherhead, P.J., Barry, F.E., Brown, G.P. & Forbes, M.R.L. (1995). Sex ratios, mating behavior and sexual size dimorphism of the northern water snake, *Nerodia sipedon*. *Behav. Ecol. Sociobiol.*, 36, 301-311.

Williams, R.N., Gopurenko, D., Kemp, K.R., Williams, B. & DeWoody, J.A. (2009). Breeding chronology, sexual dimorphism, and genetic diversity of congeneric Ambystomatid salamanders. *J. Herpetol.*, 43, 438-449.

**Table S2.** Summary of phylogenetic meta-analyses testing for a relationship between SSD and $\Delta I_s$ and $\Delta \beta_{ss}$ shown for data subsets either including an outlier (*Latrodectus hasseltii*) or excluding simultaneous hermaphrodites, which dimorphism in size is forced to be zero.

| Subset | Response | N | estimate | ± SE | $Q_M$ | *P* | $R^2$ |
|---|---|---|---|---|---|---|---|
| including *Latrodectus hasseltii* | $\Delta I_s$ | 93* | 1.014 | ± 0.324 | 9.792 | 0.002 | 0.09 |
| | $\Delta \beta_{ss}$ | 102 | 0.067 | ± 0.166 | 0.164 | 0.685 | 0.01 |
| excluding simultaneous hermaphrodites | $\Delta I_s$ | 86 | 1.048 | ± 0.334 | 9.867 | 0.002 | 0.09 |
| | $\Delta \beta_{ss}$ | 95 | 0.064 | ± 0.318 | 0.040 | 0.842 | 0.01 |

*Estimate of $\Delta I_s$ was not available for *L. hasseltii* so that results remain unaffected.

Supplementary Data

### List of primary studies

The list below encompasses all 70 published primary studies reporting data to compute lnCVR ($I_s$) and Hedges'g ($\Delta\beta_{ss}$) used in the meta-analysis. It does not include an unpublished study by Fromonteil et al. (in prep.).


Andrade, M.C.B. & Kasumovic, M.M. (2005). Terminal investment strategies and male mate choice: Extreme tests of Bateman. *Integrative and Comparative Biology*, 45, 838-847.

Anthes, N., David, P., Auld, J.R., Hoffer, J.N., Jarne, P., Koene, J.M. *et al.* (2010). Bateman gradients in hermaphrodites: an extended approach to quantify sexual selection. *Am. Nat.*, 176, 249-263.

Aronsen, T., Berglund, A., Mobley, K.B., Ratikainen, I.I. & Rosenqvist, G. (2013). Sex ratio and density affect sexual selection in a sex-role reversed fish. *Evolution*, 67, 3243-3257.

Balenger, S., Johnson, L. & Masters, B. (2009). Sexual selection in a socially monogamous bird: male color predicts paternity success in the mountain bluebird, *Sialia currucoides*. *Behav. Ecol. Sociobiol.*, 63, 403-411.

Barreto, F.S. & Avise, J.C. (2010). Quantitative measures of sexual selection reveal no evidence for sex-role reversal in a sea spider with prolonged paternal care. *Proceedings of the Royal Society B-Biological Sciences*, 277, 2951-2956.

Becher, S.A. & Magurran, A.E. (2004). Multiple mating and reproductive skew in Trinidadian guppies. *Proceedings of the Royal Society B-Biological Sciences*, 271, 1009-1014.

Bergeron, P., Montiglio, P.O., Reale, D., Humphries, M.M. & Garant, D. (2012). Bateman gradients in a promiscuous mating system. *Behav. Ecol. Sociobiol.*, 66, 1125-1130.

Bjork, A. & Pitnick, S. (2006). Intensity of sexual selection along the anisogamy-isogamy continuum. *Nature*, 441, 742-745.

Bolopo, D., Canestrari, D., Martinez, J.G., Roldan, M., Macias-Sanchez, E., Vila, M. *et al.* (2017). Flexible mating patterns in an obligate brood parasite. *Ibis*, 159, 103-112.

Borgerhoff Mulder, M. (2009). Serial monogamy as polygyny or polyandry? *Human Nature*, 20, 130-150.

Borgerhoff Mulder, M. & Ross, C.T. (2019). Unpacking mating success and testing Bateman's principles in a human population. *Proceedings of the Royal Society B-Biological Sciences*, 286, 10.

Broquet, T., Jaquiery, J. & Perrin, N. (2009). Opportunity for sexual selection and effective population size in the lek-breeding European treefrog (*Hyla arborea*) *Evolution*, 63, 674-683.

Byers, B.E., Mays, H.L., Stewart, I.R.K. & Westneat, D.F. (2004). Extrapair paternity increases variability in male reproductive success in the chestnut-slded warbler (*Dendroica pensylvanica*), a socially monogamous songbird. *Auk*, 121, 788-795.

Cattelan, S., Evans, J.P., Garcia-Gonzalez, F., Morbiato, E. & Pilastro, A. (2020). Dietary stress increases the total opportunity for sexual selection and modifies selection on condition-dependent traits. *Ecol. Lett.*, 23, 447-456.

Collet, J., Richardson, D.S., Worley, K. & Pizzari, T. (2012). Sexual selection and the differential effect of polyandry. *Proc. Natl. Acad. Sci. U. S. A.*, 109, 8641-8645.

Courtiol, A., Pettay, J.E., Jokela, M., Rotkirch, A. & Lummaa, V. (2012). Natural and sexual selection in a monogamous historical human population. *Proc. Natl. Acad. Sci. U. S. A.*, 109, 8044-8049.

Croshaw, D.A. (2010). Quantifying sexual selection: a comparison of competing indices with mating system data from a terrestrially breeding salamander. *Biol. J. Linnean Soc.*, 99, 73-83.

Devost, E. & Turgeon, J. (2016). The combined effects of pre- and post-copulatory processes are masking sexual conflict over mating rate in *Gerris buenoi*. *Journal of Evolutionary Biology*, 29, 167-177.

Fitze, P.S. & Le Galliard, J.F. (2011). Inconsistency between different measures of sexual selection. *Am. Nat.*, 178, 256-268.

Fritzsche, K. & Arnqvist, G. (2013). Homage to Bateman: sex roles predict sex differences in sexual selection. *Evolution*, 67, 1926-1936.

Fuxjager, L., Wanzenbock, S., Ringler, E., Wegner, K.M., Ahnelt, H. & Shama, L.N.S. (2019). Within-generation and transgenerational plasticity of mate choice in oceanic stickleback under climate change. *Philos. Trans. R. Soc. B-Biol. Sci.*, 374, 12.

Gagnon, M.-C., Duchesne, P. & Turgeon, J. (2012). Sexual conflict in *Gerris gillettei* (Insecta: Hemiptera): influence of effective mating rate and morphology on reproductive success. *Canadian Journal of Zoology*, 90, 1297-1306.

Garcia-Navas, V., Ferrer, E.S., Bueno-Enciso, J., Barrientos, R., Sanz, J.J. & Ortego, J. (2014). Extrapair paternity in Mediterranean blue tits: socioecological factors and the opportunity for sexual selection. *Behavioral Ecology*, 25, 228-238.

Gerlach, N.M., McGlothlin, J.W., Parker, P.G. & Ketterson, E.D. (2012). Reinterpreting Bateman gradients: multiple mating and selection in both sexes of a songbird species. *Behavioral Ecology*, 23, 1078-1088.

Glaudas, X., Rice, S.E., Clark, R.W. & Alexander, G.J. (2020). The intensity of sexual selection, body size and reproductive success in a mating system with male-male combat: is bigger better? *Oikos*, 129, 998-1011.

Gopurenko, D., Williams, R.N. & DeWoody, J.A. (2007). Reproductive and mating success in the small-mouthed salamander (*Ambystoma texanum*) estimated via microsatellite parentage analysis. *Evolutionary Biology*, 34, 130-139.

Gopurenko, D., Williams, R.N., McCormick, C.R. & DeWoody, J.A. (2006). Insights into the mating habits of the tiger salamander (*Ambystoma tigrinum tigrinum*) as revealed by genetic parentage analyses. *Mol. Ecol.*, 15, 1917-1928.

Grunst, A.S., Grunst, M.L., Korody, M.L., Forrette, L.M., Gonser, R.A. & Tuttle, E.M. (2019). Extrapair mating and the strength of sexual selection: insights from a polymorphic species. *Behavioral Ecology*, 30, 278-290.

Hoffer, J.N.A., Marien, J., Ellers, J. & Koene, J.M. (2017). Sexual selection gradients change over time in a simultaneous hermaphrodite. *eLife*, 6, 16.

Janicke, T., David, P. & Chapuis, E. (2015). Environment-dependent sexual selection: Bateman's parameters under varying levels of food availability. *Am. Nat.*, 185, 756-768.

Jokela, M., Rotkirch, A., Rickard, I.J., Pettay, J. & Lummaa, V. (2010). Serial monogamy increases reproductive success in men but not in women. *Behavioral Ecology*, 21, 906-912.

Jones, A.G., Arguello, J.R. & Arnold, S.J. (2002). Validation of Bateman's principles: a genetic study of sexual selection and mating patterns in the rough-skinned newt. *Proc. R. Soc. Lond. Ser. B-Biol. Sci.*, 269, 2533-2539.

Jones, A.G., Arguello, J.R. & Arnold, S.J. (2004). Molecular parentage analysis in experimental newt populations: The response of mating system measures to variation in the operational sex ratio. *Am. Nat.*, 164, 444-456.

Jones, A.G., Rosenqvist, G., Berglund, A., Arnold, S.J. & Avise, J.C. (2000). The Bateman gradient and the cause of sexual selection in a sex-role-reversed pipefish. *Proc. R. Soc. Lond. Ser. B-Biol. Sci.*, 267, 677-680.

Jones, P.H., Van Zant, J.L. & Dobson, F.S. (2012). Variation in reproductive success of male and female Columbian ground squirrels (*Urocitellus columbianus*). *Can. J. Zool.-Rev. Can. Zool.*, 90, 736-743.

Ketterson, E.D., Parker, P.G., Raouf, S.A., Nolan Jr, V., Ziegenfus, C. & Chandler, C.H. (1997). The relative impact of extra-pair fertilizations on variation in male and female reproductive success in dark-eyed juncos (*Junco hyemaus*). In: *Avian Reproductive Tactics: Female and Male Perspectives* (eds. Parker, PG & Burley, NT), pp. 81-101.

Krakauer, A.H. (2008). Sexual selection and the genetic mating system of Wild Turkeys. *Condor*, 110, 1-12.

Kretzschmar, P., Auld, H., Boag, P., Ganslosser, U., Scott, C., de Groot, P.J.V. *et al.* (2020). Mate choice, reproductive success and inbreeding in white rhinoceros: New insights for conservation management. *Evol. Appl.*, 13, 699-714.

Levine, B.A., Schuett, G.W., Clark, R.W., Repp, R.A., Herrmann, H.W. & Booth, W. (2020). No evidence of male-biased sexual selection in a snake with conventional Darwinian sex roles. *R. Soc. Open Sci.*, 7, 10.

Levine, B.A., Smith, C.F., Schuett, G.W., Douglas, M.R., Davis, M.A. & Douglas, M.E. (2015). Bateman-Trivers in the 21st Century: sexual selection in a North American pitviper. *Biol. J. Linnean Soc.*, 114, 436-445.

Levitan, D.R. (2008). Gamete traits influence the variance in reproductive success, the intensity of sexual selection, and the outcome of sexual conflict among congeneric sea urchins. *Evolution*, 62, 1305-1316.

Louder, M.I.M., Hauber, M.E., Louder, A.N.A., Hoover, J.P. & Schelsky, W.M. (2019). Greater opportunities for sexual selection in male than in female obligate brood parasitic birds. *Journal of Evolutionary Biology*, 32, 1310-1315.

Mangold, A., Trenkwalder, K., Ringler, M., Hoedl, W. & Ringler, E. (2015). Low reproductive skew despite high male-biased operational sex ratio in a glass frog with paternal care. *Bmc Evolutionary Biology*, 15.

Marie-Orleach, L., Janicke, T., Vizoso, D.B., David, P. & Scharer, L. (2016). Quantifying episodes of sexual selection: Insights from a transparent worm with fluorescent sperm. *Evolution*, 70, 314-328.

McCullough, E.L., Buzatto, B.A. & Simmons, L.W. (2018). Population density mediates the interaction between pre- and postmating sexual selection. *Evolution*, 72, 893-905.

Mills, S.C., Grapputo, A., Koskela, E. & Mappes, T. (2007). Quantitative measure of sexual selection with respect to the operational sex ratio: a comparison of selection indices. *Proceedings of the Royal Society B-Biological Sciences*, 274, 143-150.

Mobley, K.B. & Jones, A.G. (2013). Overcoming statistical bias to estimate genetic mating systems in open populations: a comparison of Bateman's principles between the sexes in a sex-role-reversed pipefish. *Evolution*, 67, 646-660.

Moorad, J.A., Promislow, D.E.L., Smith, K.R. & Wade, M.J. (2011). Mating system change reduces the strength of sexual selection in an American frontier population of the 19th century. *Evolution and Human Behavior*, 32, 147-155.

Morimoto, J., Pizzari, T. & Wigby, S. (2016). Developmental environment effects on sexual selection in male and female *Drosophila melanogaster*. *PLoS One*, 11, 27.

Munroe, K.E. & Koprowski, J.L. (2011). Sociality, Bateman's gradients, and the polygynandrous genetic mating system of round-tailed ground squirrels (*Xerospermophilus tereticaudus*). *Behav. Ecol. Sociobiol.*, 65, 1811-1824.

Pelissie, B., Jarne, P. & David, P. (2012). Sexual selection without sexual dimorphism: Bateman gradients in a simultaneous hermaphrodite. *Evolution*, 66, 66-81.

Poesel, A., Gibbs, H.L. & Nelson, D.A. (2011). Extrapair fertilizations and the potential for sexual selection in a socially monogamous songbird. *Auk*, 128, 770-776.

Pongratz, N. & Michiels, N.K. (2003). High multiple paternity and low last-male sperm precedence in a hermaphroditic planarian flatworm: consequences for reciprocity patterns. *Mol. Ecol.*, 12, 1425-1433.

Prosser, M.R., Weatherhead, P.J., Gibbs, H.L. & Brown, G.P. (2002). Genetic analysis of the mating system and opportunity for sexual selection in northern water snakes (*Nerodia sipedon*). *Behavioral Ecology*, 13, 800-807.

Rios-Cardenas, O. (2005). Patterns of parental investment and sexual selection in teleost fishes: Do they support Bateman's principles? *Integrative and Comparative Biology*, 45, 885-894.

Rodriguez-Munoz, R., Bretman, A., Slate, J., Walling, C.A. & Tregenza, T. (2010). Natural and sexual selection in a wild insect population. *Science*, 328, 1269-1272.

Saunders, K.M. & Shuster, S.M. (2019). Bateman gradients and alternative mating strategies in a marine isopod. *IntechOpen*, (DOI: 10.5772/intechopen.88956).

Schlicht, E. & Kempenaers, B. (2013). Effects of social and extra-pair mating on sexual selection in blue tits (*Cyanistes caeruleus*) *Evolution*, 67, 1420-1434.

Schulte-Hostedde, A.I., Millar, J.S. & Gibbs, H.L. (2004). Sexual selection and mating patterns in a mammal with female-biased sexual size dimorphism. *Behavioral Ecology*, 15, 351-356.

Serbezov, D., Bernatchez, L., Olsen, E.M. & Vollestad, L.A. (2010). Mating patterns and determinants of individual reproductive success in brown trout (*Salmo trutta*) revealed by parentage analysis of an entire stream living population. *Mol. Ecol.*, 19, 3193-3205.

Skjaervo, G.R. & Roskaft, E. (2015). Wealth and the opportunity for sexual selection in men and women. *Behavioral Ecology*, 26, 444-451.

Tatarenkov, A., Healey, C.I.M., Grether, G.F. & Avise, J.C. (2008). Pronounced reproductive skew in a natural population of green swordtails, *Xiphophorus helleri*. *Mol. Ecol.*, 17, 4522-4534.

Ursprung, E., Ringler, M., Jehle, R. & Hodl, W. (2011). Strong male/male competition allows for nonchoosy females: high levels of polygynandry in a territorial frog with paternal care. *Mol. Ecol.*, 20, 1759-1771.

Walker, L.K., Ewen, J.G., Brekke, P. & Kilner, R.M. (2014). Sexually selected dichromatism in the hihi *Notiomystis cincta*: multiple colours for multiple receivers. *Journal of Evolutionary Biology*, 27, 1522-1535.

Wang, X., Liu, S., Yang, Y.Q., Wu, L.N., Huang, W.H., Wu, R.X. *et al.* (2020). Genetic evidence for the mating system and reproductive success of black sea bream (*Acanthopagrus schlegelii*). *Ecol. Evol.*, 10, 4483-4494.

Wells, C.P., Tomalty, K.M., Floyd, C.H., McElreath, M.B., May, B.P. & Van Vuren, D.H. (2017). Determinants of multiple paternity in a fluctuating population of ground squirrels. *Behav. Ecol. Sociobiol.*, 71, 13.

Whittingham, L.A. & Dunn, P.O. (2005). Effects of extra-pair and within-pair reproductive success on the opportunity for selection in birds. *Behavioral Ecology*, 16, 138-144.

Whittingham, L.A. & Lifjeld, J.T. (1995). High paternal investment in unrelated young: extra-pair paternity and male parental care in house martins. *Behav. Ecol. Sociobiol.*, 37, 103-108.

Williams, R.N. & DeWoody, J.A. (2009). Reproductive success and sexual selection in wild eastern tiger salamanders (*Ambystoma t. tigrinum*). *Evolutionary Biology*, 36, 201-213.

Woolfenden, B.E., Gibbs, H.L. & Sealy, S.G. (2002). High opportunity for sexual selection in both sexes of an obligate brood parasitic bird, the brown-headed cowbird (*Molothrus ater*). *Behav. Ecol. Sociobiol.*, 52, 417-425.
